# Detection of pre-microRNA with Convolutional Neural Networks

**DOI:** 10.1101/840579

**Authors:** Jorge Cordero, Vlado Menkovski, Jens Allmer

**Affiliations:** Department of Mathematics and Computer Science, Eindhoven University of Technology, Eindhoven, The Netherlands; Medical Informatics an Bioinformatics, Hochschule Ruhr West, University of Applied Sciences, Mülheim an der Ruhr, Germany

## Abstract

MicroRNAs (miRNAs) are small non-coding RNA sequences that have been implicated in many physiological processes and diseases. The experimental discovery of miRNAs is complicated because both miRNAs and their targets need to be expressed for the confirmation of functional interactions, but expression is under spatiotemporal control. This has motivated the development of computational methods for miRNA detection. This typically involves feature design by domain experts followed by machine learning. While handcrafted features can encode domain knowledge, feature engineering is a time-consuming task. Additionally, some of the currently most successful features for pre-miRNA detection, such as p-value based ones, require comparably large computations. In contrast, advances of representation learning methods such as deep learning can discover relevant features directly from data. Here, we propose a method that uses domain knowledge to create an efficient graphical representation of pre-miRNAs, encoding sequence, structure, and implicitly some thermodynamic information. A suitable convolutional neural network architecture for pre-miRNA detection was used to train a model. This model achieves state-of-the-art performance on all previously used datasets. Additionally, computations succeed in real time thereby overcoming current speed limitations. Finally, our strategy promises future interpretability of the trained models and in turn novel biological interpretations of pre-miRNA characteristics.

## 1. Introduction

Mature microRNAs (miRNAs) are small noncoding RNAs 18-24 nucleotides long involved in regulating gene expression. They derive from longer RNA sequences called precursor miRNAs (pre-miRNAs) via a controlled pathway (1). These miRNAs are involved in most biological processes, ranging from cell differentiation, organ development, and angionesis to apoptosis (2, 3). In humans, deregulated miRNAs have been associated with cancer (4, 5), autoimmune diseases (6), and neurological disorders (5). Due to their presence in body fluids, miRNAs are considered good candidates for noninvasive biomarkers. As of today, hundreds of miRNAs have been identified for several species and many more are predicted to exist.

There are many methods to experimentally detect miRNAs, but they are quite involved and can only detect actually expressed miRNAs. However, many miRNAs and their targets that are only expressed under specific conditions, therefore, escape detection. This has led to the development of computational methods to aid the detection of miRNAs and pre-miRNAs. Among these methods, machine learning (ML) algorithms have become widely used for pre-miRNA detection (7). These algorithms train models using features usually derived from primary sequence, secondary structure, and thermodynamic properties, or mixtures thereof, as well as mathematical transformations. The trained models are then used to detect pre-miRNAs from candidate sequences.

ML approaches rely on engineered features created by domain experts to produce quality results; however, developing relevant features is time consuming and error prone as it has major impact on the performance of ML models (8). Nonetheless, an abundance of features has been described to parameterize pre-miRNAs (9). For instance, dinucleotide frequencies (10) quantify primary sequence composition, local contiguous triplet elements (11) measure primary sequence composition and secondary structure combined, and thermodynamic properties such as enthalpy (12) quantify the stability of the secondary structure. While these features allow ML models to learn useful parameters for classification, they also restrict the amount of useful information models can access. Thus, ML practitioners frequently perform feature selection procedures to select the most informative features to train useful models. The effects of feature selection in pre-miRNA prediction have been shown in the works of (13, 14), where a large number of features are reduced to the most informative ones. Furthermore, the performance of ML models is also tied to the training and testing datasets. This effect is reported in a study of all *ab initio* pre-miRNA classification methods published previously (15).

In this work, we aim to build a pre-miRNA detection procedure that uses deep learning (DL) models. DL models automatically learn high level informative features that are used to compute the model’s output by applying a series of transformations to low level weakly informative features corresponding to the model’s input. DL has enabled solutions to many long-standing ML problems in domains such as image analysis, natural language processing, and speech recognition. Following the success of convolutional neural networks (CNNs) on image classification challenges, such as the AlexNet (16) and ResNet (17), these DL methods have been recently applied in bioinformatics. For instance, CNNs have been used for gene expression prediction (18) and DNA sequence binding prediction (19). Most notably, DeepVariant (20) encodes sequence alignments as images that are used to train CNN models for the prediction of genetic variation.

Regarding pre-miRNA prediction, the work of Do et al. (21) employs deep CNNs to identify pre-miRNAs from candidate sequences encoded as matrices containing information of primary sequence, secondary structure, and minimum free energy. The latter, although using DL, relies on hand crafted features, encoding the information into what they call pairing matrices as input for their 2D CNN.

Here, we leverage the fact that CNNs can capture localized spatial patterns and infer features based on these patterns. We propose a pre-miRNA detection framework (Figure 1) that consists of an encoding procedure that generates intermediate representations of nucleotide sequences and a CNN model that uses these representations for pre-miRNA detection. The encoding procedure (Figure 2) transforms significant characteristics of the primary sequence and secondary structure into color images that expose potential patterns in the pre-miRNA structure. These images allow domain experts to visually inspect relevant characteristics of nucleotide sequences, but more importantly, they can be used by CNNs to automatically build useful higher level features for pre-miRNA detection. Additionally, this encoding allows us to interpret the decisions made by CNNs via inspecting the corresponding saliency maps. These maps show the effect that each region (pixel) of an image has on the decision of a CNN model allowing for the interpretation of the relationship between input, image, and model decision.

**Fig. 1.**
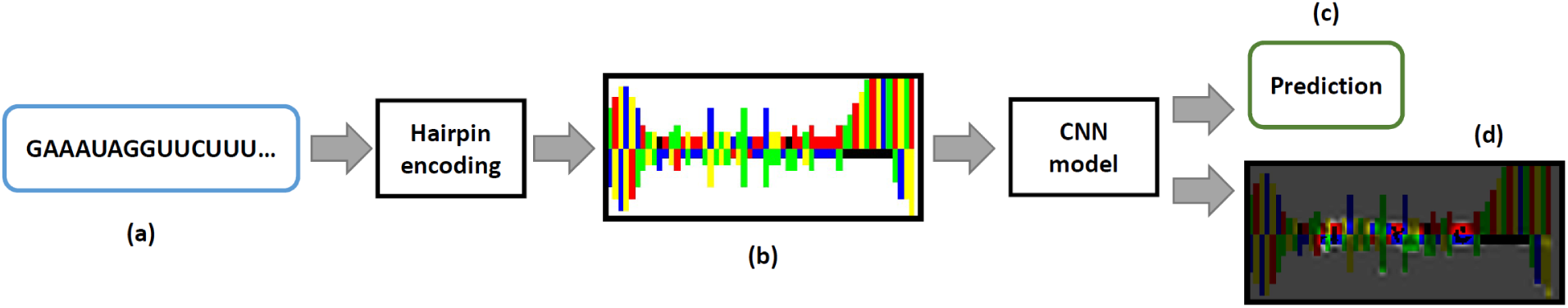
Precursor miRNA detection framework. First, a nucleotide sequence (a) is encoded into an RGB image (b) that represents its hairpin structure. Then, using a CNN model, the corresponding label (c) is predicted. Additionally, a saliency map (d) can be computed to explore the pixels that influence the prediction.

**Fig. 2.**
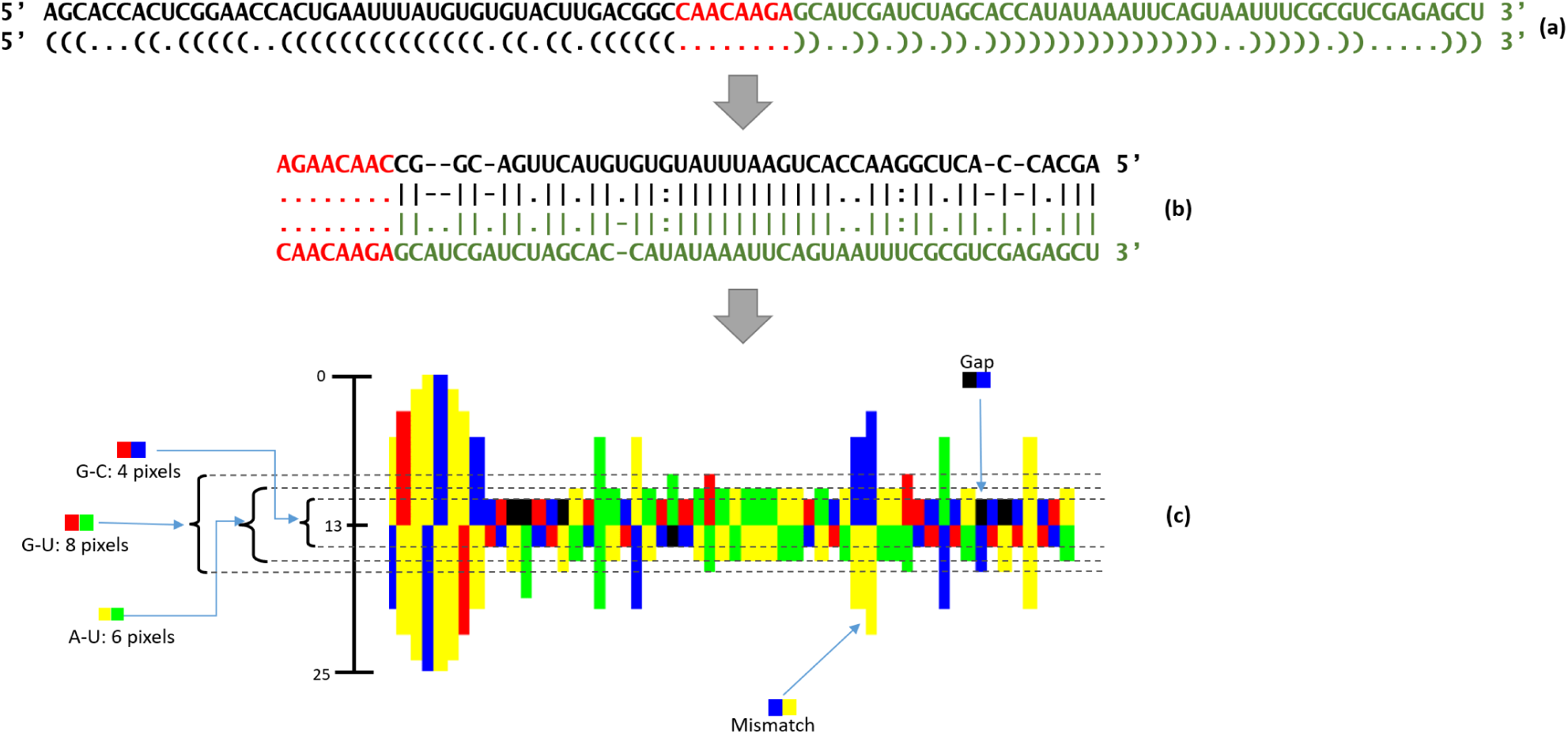
Encoding the cel-mir-83 pre-miRNA as color image. From a primary RNA sequence, the secondary structure is computed using RNAFold with default values (a). The structure is presented in the dot bracket format underneath the sequence. The computed fold and primary sequence are then co-aligned, with the hairpin section (b). The terminal loop (red section) is included twice and is left aligned. Finally, the corresponding color image is generated (c). G-C bonds are represented with four pixels centered around the middle (pixel 13); two pixels per nucleotide. Similarly, A-U bonds are represented with six pixels, and G-U bonds are represented with eight pixels. Notice that mismatches and gaps can have different lengths according to their neighbors.

We implemented our approach in the tool DeepMir, which is available at https://github.com/jacordero/deepmir. Application of DeepMir to datasets used by Saçar et al. (15) revealed that our framework identifies pre-miRNAs of several species better than SVMs, Decision Trees, Naive Bayes models, as well as ensemble models trained using different sets of features. More importantly, DeepMir outperforms the best ML models in this work (mostly ensemble models) at classifying candidate sequences from novel datasets. The efficiency and accuracy of DeepMir for pre-miRNA detection makes it applicable to the analysis of large eukaryotic genomes.

## 2. Materials and Methods

### Datasets

Nucleotide sequences from various positive and negative datasets, taken from a study by Saçar et al. (15), were encoded as hairpin images to train and evaluate CNN models:

- miRBase: pre-miRNAs downloaded from the miRBase 21st release (http://www.mirbase.com) containing 28,596 examples from several species
- hsa: subset of miRBase containing 1,881 human pre-miRNAs
- mmu: subset of miRBase containing 1,193 mouse pre-miRNAs
- mmu*: subset of 380 mouse pre-miRNAs extracted from miRBase with at least 100 reads per million
- MirGeneDB: set of 1,434 pre-miRNAs downloaded from http://www.mirgenedb.org
- hsa+: subset of 523 human pre-miRNAs extracted from MirGeneDB
- mmu+: subset of 395 mouse pre-miRNAs extracted from MirGeneDB
- gga+: subset of 229 chicken pre-miRNAs extracted from MirGeneDB
- dre+: subset of 287 zebra fish pre-miRNAs extracted from MirGeneDB
- NegHsa: negative dataset containing 68,046 unique sequences introduced by Gudys et al. (22)
- Pseudo: negative dataset composed of 8,492 pseudo pre-miRNAs proposed by Ng et al. (10)
- Shuffled: negative dataset containing 1,423 entries created by shuffling human pre-miRNAs
- Zou: negative sequences generated by applying a sample selection technique to sub-sequences obtained from coding regions of known mature miRNAs (23)
- Chen: negative examples sampled from Zou and Pseudo (24)
- NotBestFold: negative dataset containing suboptimal folds of pre-miRNAs from miRBase (15)

Using these positive and negative example datasets, we created two datasets to train CNN models for pre-miRNA detection, *modmiRBase* and *modhsa*. The *modmiRBase* dataset contains positive examples obtained from miRBase after removing duplicate sequences. The *modhsa* dataset contains only the unique human hairpin images present in *modmiRBase*. Negative examples for both datasets were obtained from the Pseudo, Shuffled, and NegHsa datasets.

Joining the positive and negative elements directly would result in heavily unbalanced datasets. For instance, the *modhsa* dataset would have a negative to positive ratio of about 40:1 (3:1 for *modmiRBase*). For such an unbalanced dataset, it is easy to obtain high accuracy values on classification tasks by building classifiers that output the majority class label. However, the recall values are typically very low for such classifiers. To reduce the effects that unbalanced datasets can have on trained classifiers, Weis (25) recommends techniques to overcome issues with unbalanced datasets such as sampling methods. Accordingly, in this work, we use a subsampling method to create datasets having the same number of positive and negative entries. For the *modhsa* dataset, the same number of negative examples are selected in a stratified manner from each of the three selected negative datasets. In contrast, for the *modmiRBase* dataset, we include all elements from Pseudo and Shuffled but sampled a subset of images from NegHsa.

### Hairpin Encoding

In our approach we aim to leverage the power of deep CNN models to develop effective representations of the low-level high dimensional data that enable accurate classification. On the other hand, we also would like to benefit from available knowledge on the base pair interaction and their bond strengths (thermodynamic information) as these properties play a major role in the forming of miRNA molecules. We, therefore, developed an encoding of the pre-miRNA sequence information as an image, enriched with its secondary structure and implicit thermodynamic information.

For a given nucleotide sequence, the encoding procedure works as follows. First, the secondary structure is predicted using RNAFold (26) with default settings (see Figure 2(a)). Then, as illustrated in Figure 2(b), the resulting fold and sequence are aligned according to the structural constraints given by the parentheses in the predicted secondary structure with the complete terminal loop (depicted in red) duplicated on the left. Next, we represent this structure as an image by using different colors to encode nucleotides and gaps (A: yellow, C: blue, G: red, U: green, and gap: black). The strength of the bonds between the nucleotides are represented by extending the nucleotide pixels from the middle up and down towards the top and bottom of the image accordingly. Specifically, the length of the bar representing the nucleotide pairs takes into account different bond strengths: G-C > A-U > G-U > gap. G-C bonds are represented with two pixels above and below the mid-line of the image using the appropriate colors defined above. A-U bonds get an additional pixel and G-U wobbles two pixels, in order to represent lower bond strengths compared to G-C bonds, towards the top and bottom of the image, respectively. The final strength of the bond is not only defined by the nucleotide pair, but by their neighbouring pairs as well. This spatial context is represented by extending the nucleotide pixels (weakening the bond) when a pair is adjacent to other mismatches or gaps. So, a sequence of consecutive mismatches is represented by an additional extension of the nucleotide pixels in the middle of the section (see Figure 2(c)). Finally, this representation is pasted on a fixed sized image canvas of 100×25 pixels. This results in pre-miRNAs larger than 100 base pairs being cleaved on the stem side, which is an acceptable compromise since the mature miRNA is expected to be close to the loop and typically consist of around 18-24 nucleotides.

With respect to time, the hairpin encoding algorithm encodes sequences efficiently. A runtime analysis shows that RNAFold dominates the running time of the encoding procedure. For a given nucleotide sequence of length *m*, RNAFold takes *O*(*m*^3^), and the alignment is a trivial process that is hardly measurable. A simple implementation of the drawing procedure uses two loops and runs in *O*(*m*^2^). An outer loop draws each nucleotide according to the predefined bond strengths, and an inner loop extends each nucleotide pixel inside a sequence of mismatches according to its position in the sequence. Compared to feature extraction methods, our algorithm allows faster encoding of pre-miRNA candidates from a genome. For instance, as parsing the human genome takes *O*(*n*), with *n* being the length of the genome, encoding human pre-miRNA candidates requires *O*(*nm*^3^).

### CNN Models

Many successful CNN architectures have been proposed for natural image processing. However, the synthetic images generated by our hairpin encoding method have different spatial patterns than the data used to train these CNN models. Therefore, we developed custom based architectures that use architectural ideas and components from successful models such as Visual Geometry Group (VGG) (27), GoogLeNet (28), and ResNet (17). We tested a number of models (48) that include different variations of successful practices in CNN design. The architecture of these models can be found in Tables 3 to 5, and the performance of each trained model is shown in Tables 7 to 9.

Among all designed CNNs, the model named VGG-Style-3M-256FU obtained the best performance. This CNN (depicted in Figure 3) is based on the VGG architecture and contains three convolutional modules and one fully connected module. The convolutional modules have two convolutional layers each, where each convolutional layer has a fixed number of filters (48, 60, or 72) with shape 3×3. Then, a max-pool layer with filter shape 2×2 and a dropout layer follow the convolutional layers. Finally, the fully connected module maps the features obtained in the last convolutional module to the output that represent the probability of the input molecule being detected as pre-miRNA. This module is composed by a fully connected layer with 256 units, a dropout layer, and a softmax output layer.

**Fig. 3.**
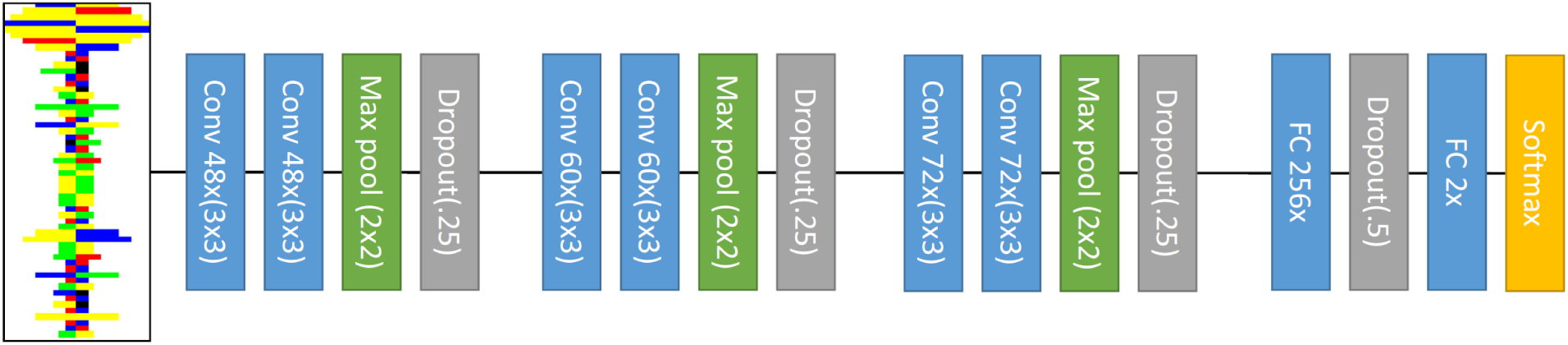
Elements of the VGG-Style-3M-256FU network. For the convolutional layers, the parameters describe number of filters and filter size respectively. For the maxpool layers, the parameters describe both filter size and stride values. The fully connected layers indicate their number of units.

### Saliency maps for pre-miRNA images

The image representation of the pre-miRNA and the CNN model enable us to visualize the salient regions (pixels relevant for the decision making of the model) of those images. These salient regions indicate to which parts of the molecule our model is sensitive. We obtain saliency maps using smoothed guided backpropagation (29, 30). Sensitivity analysis such as this allows for interpretations of the behaviour of the CNN model. As DL models and other complex machine learning models are typically black-boxes, enabling interpretation poses a significant advantage for the expert evaluation of their decisions which in turn results in higher adoption rates of the model.

## 3. Experiments and Results

Using a set of 1,000 randomly selected sequences from miRBase, we evaluated the performance of the hairpin encoding algorithm in two ways: measuring the time required to create a set of images and measuring the time of each step required to create an image. For these measurements, the sequences were loaded to the RAM memory of a PC with an Intel i7 processor at 3.3GHz and 32GB of RAM. In the first evaluation, for each sequence, we computed a hairpin image and the features described in Saçar et al. According to the results in Table 10, on average, our image encoding algorithm takes nine seconds while computing features requires about six hours. Notice that creating hairpin images is more than 2,000 times faster than computing hundreds of features. For large genomes, these results represent a significant improvement in the computation of pre-miRNA representations. The values in Table 11 show that RNAFold is the most time-consuming step of our hairpin encoding algorithm. We observe that, on average, folding takes about 95% of the total time required to encode a single sequence. For instance, RNAFold takes 29 milliseconds (minimum of 16ms; maximum of 314ms at a length of 500nt) while the alignment, drawing, and file writing steps take 1.15 milliseconds combined. These results agree with the runtime analysis presented in the previous section.

For the training of DL models with the architectures outlined in Figure 3 and Tables 3 to 5 positive and negative examples of pre-miRNAs are needed. As positive examples, we selected 1,830 human hairpin images and combined them with 1,830 negative example images taken from the Pseudo, Shuffled, and NegHsa datasets (610 examples from each dataset). In the following, we will refer to this dataset as the *modhsa* dataset. Since this dataset is limited in size, but the largest available for a single species, we also develop a second, larger dataset that contains 24,801 images containing all species in miRBase as positive examples and an equal number of negative images selected from the Pseudo, Shuffled, and NegHsa datasets. We will refer to this larger dataset as *modmiRBase* in the following.

Initially, CNN models were trained using 70% of the hairpin images randomly selected from *modhsa*, while the remaining hairpin images were used for testing. These models were implemented using the Keras library (31). For training, we used the Adam (32) optimization algorithm with default values (learning rate of 0.001, *β*^1^ = 0.9, *β*^2^ = 0.999, *ϵ* = 1^−7^, and decay of 0), 100 epochs as the number of iterations over the entire training dataset, and a batch size of 128 corresponding to the number of samples used by Adam to update the CNN parameters. Training directly on the *modhsa* dataset, we achieve ∼ 92% accuracy.

The *modhsa* dataset can be considered small for training deep learning models, as they tend to perform better when trained on large datasets. For some image analysis tasks, such as digit recognition or object recognition, it is relatively easy to collect more positive examples. In contrast, in this study we were limited to previously verified pre-miRNAs. Additionally, data augmentation techniques typically used in image analysis where transformation of the image data is possible without changing the target classification are also not suitable for this case. We did, however, in our case benefit from using a transfer learning approach, i.e., we pre-trained the CNNs on different datasets and then fine-tuned the models on the *modhsa* dataset. Transfer learning typically provides advantages when restricted amounts of data are available. The strategy is to develop good low-level detectors on data with similar properties (or solving a similar task), which can then effectively be used on the target task to boot-strap the training process.

Specifically, we pre-trained the model using a subset of hairpin images from *modmiRBase* that does not contain images in *modhsa*. These subset of images contains a larger number of pre-miRNAs that share similar statistical properties to human pre-miRNAs such as the distribution of minimum free energy of folding and sequence length distribution. The pre-training procedure was performed using 40 epochs, Adam with default values, and a batch size of 128. The resulting pre-trained models were then fine-tuned using 70% of the entries in the *modhsa* dataset, and the remaining 30% were used for testing. For fine-tuning, we used 100 epochs, Adam with default values, and the same batch size.

As shown in Tables 7 to 9, the CNN models resulting from pre-training outperform the models without pre-training. Overall, the best pre-trained CNN reaches 95% accuracy while the best non pre-trained CNN only reaches 92%. The performance of our best pre-trained models and their corresponding non pre-trained versions is displayed in Table 6. From these results, we select the pre-trained VGG-Style-3M-256FU network as our best CNN model (fine-tuned-CNN).

We compare the performance of our models with existing work employing SVM, decision trees (DTs), and Naive Bayes (NB) models trained by Saçar et al. (15) using features defined in the works of Xue (11), Ng (10), Batuwita (12), and Gudys (22), among others. The balanced dataset used to train these models contains human hairpins obtained from miRBase and negative sequences sampled from the Pseudo dataset. The dataset was split 70%-30% for training and testing respectively. In the study by Saçar et al. the average performance over 1000fold MCCV for SVM, DT, and NB models was computed for varying feature sets. Here, features are automatically learned from the graphical representation. We here evaluated a large number of models by training and testing CNNs derived from different architectures and picked a model in that process. Since many models are not achieving good performance and because this is not an inherent discriminator for the other models, we present the best model instead of the average model accuracy. The accuracy values of the selected ML models, our best CNN, and the corresponding CNN without pre-training (base-CNN), are shown in Table 1. We observe a considerable difference in performance between the best ML model (Chen) and the fine-tuned-CNN model. These results suggest that CNNs can learn to extract relevant features from the hairpin images and develop accurate detection model for pre-miRNAs.

**Table 1.** CNN models compared against the best ML models selected among trained instances of DTs, NBs, and SVMs trained by Saçar et al. These ML models were trained and tested using human pre-miRNAs from miRBase and negative sequences from the Pseudo dataset. The base-CNN model was trained using only the *modhsa* dataset and the fine-tuned-CNN model was pre-trained using *modhsa* and fine-tuned using *modmiRBase*.

| Model | Number of features | Accuracy |
| --- | --- | --- |
| fine-tuned-CNN | 2 | **0.952** |
| base-CNN | 2 | 0.917 |
| Chen | 99 | 0.913 |
| Ng | 29 | 0.899 |
| Ding | 32 | 0.888 |
| Batuwita | 21 | 0.875 |
| Gudys | 28 | 0.874 |
| Burgt | 18 | 0.872 |
| Bentwich | 26 | 0.870 |
| Gao | 57 | 0.863 |
| Jiang | 34 | 0.862 |
| Lopes | 13 | 0.860 |
| Xu | 35 | 0.857 |
| Ritchie | 36 | 0.849 |
| Xue | 32 | 0.811 |

ROC curves computed for both CNNs (see Figure 4) show that fine-tuned-CNN excels at discriminating between positive and negative examples. Hence, in case we need to trade accuracy to reduce the false positive rate, this model could still perform similar to the best ML models from Table 1. Additionally, the ROC curves also indicate that our models excel at identifying negative examples, which is expected because both models were trained using balanced datasets.

**Fig. 4.**
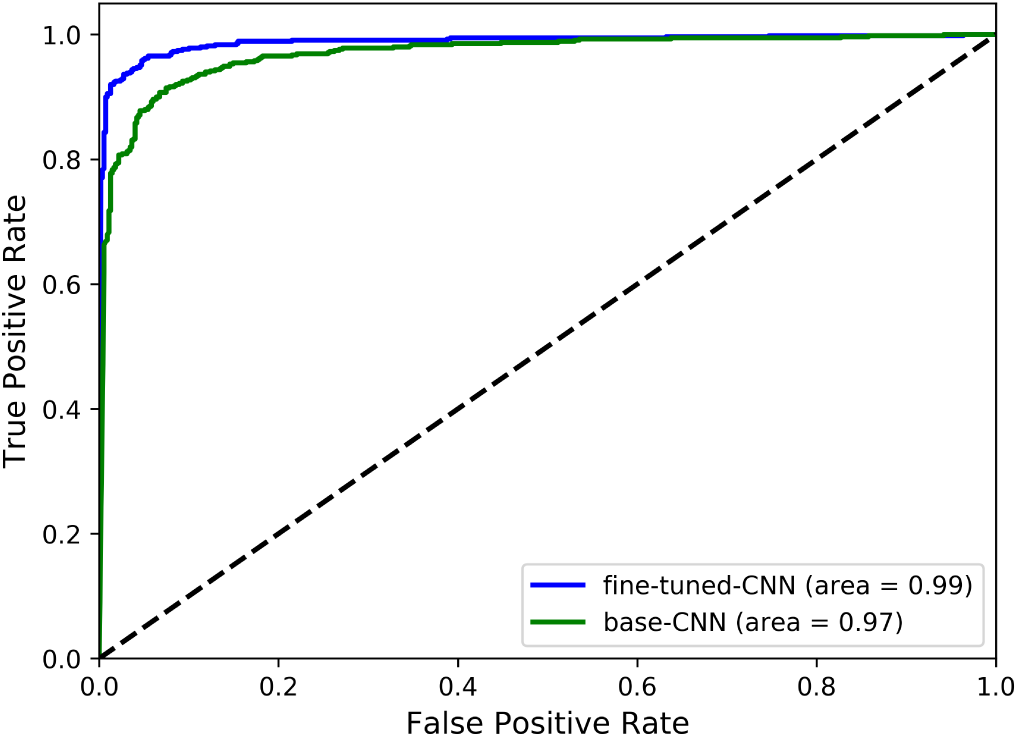
ROC curves for fine-tuned-CNN and base-CNN. The area under the curve for both models shows they have an excellent performance predicting positive and negative pre-miRNAs.

Models for pre-miRNA detection are expected to identify novel pre-miRNAs. Due to the nature of our study, we could not validate that our framework can detect unknown pre-miRNAs using experimental methods as done for example by Bentwich and colleagues (33). Instead, as done in most studies concerned with pre-miRNA prediction (e.g.:, (14, 34)), we used several positive and negative datasets to observe the performance of our CNNs when applied to previously unseen hairpin images. We consider that if our models perform well at classifying unseen nucleotide sequences, they should perform equally well with novel pre-miRNAs. Therefore, we evaluated our models using several positive and negative datasets and compared their performance against the best two ensemble models employed by Saçar et al. (Average_*DT*_ and Consensus_*NB*_). The results shown in Table 2 indicate that the four models are better at predicting positive sequences than negative ones. For instance, all models obtain a true negative rate lower than 0.6 on the Zou dataset (the most difficult negative dataset for both CNNs, but also for all ML approaches). On the contrary, although mmu appears to be a difficult positive dataset for both ML models, the CNNs perform well on this dataset. Overall, our CNNs obtain an excellent performance on positive and negative datasets. By looking at the average accuracy, we observe that both CNNs perform better than existing ensemble-based models. Saçar et al. previously noted, that with higher data quality (mmu < mmu* < mmu+) true positive and true negative prediction rates improve (15) and the same can be observed for CNN performance.

**Table 2.** Successful detection of non pre-miRNA for negative datasets and pre-miRNA for positive datasets. Average_*DT*_ and Consensus_*NB*_ are the best two ensemble models from Saçar et al. The fine-tuned-CNN and base-CNN models correspond to the VGG-Style-3M-256FU network with and without pre-training respectively.

| Model | Negative datasets |  |  |  |  |  | Positive datasets |  |  |  |  |  |  |  |  |
| --- | --- | --- | --- | --- | --- | --- | --- | --- | --- | --- | --- | --- | --- | --- | --- |
|  | *NegHsa* | *Zou* | *Pseudo* | *NotBestFold* | *Shuffled* | *Chen* | *hsa* | *mmu* | *mmu\** | *miRBase* | *MirGeneDB* | *hsa+* | *mmu+* | *gga+* | *dre+* |
| fine-tuned-CNN | **97** | 26 | 91 | **97** | **99** | 65 | **98** | **93** | **97** | **93** | **100** | **100** | **99** | **100** | **100** |
| base-CNN | 93 | 43 | 74 | **97** | 96 | 62 | **98** | 91 | 95 | 82 | 99 | 99 | 98 | 99 | 99 |
| $Average_{DT}$ | 82 | **56** | **93** | 31 | 93 | **77** | 97 | 83 | 95 | 91 | 97 | 98 | 96 | 98 | 96 |
| $Consensus_{NB}$ | 89 | 52 | 86 | 24 | 96 | **77** | 86 | 82 | 93 | 89 | 96 | 96 | 93 | 98 | 97 |

To further evaluate our CNN model, we selected the organisms having at least 200 entries in the complete miRBase dataset, and used these examples to compute true positive rate (TPR) values per organism. The resulting TPR values for fine-tuned-CNN and both ensemble models are depicted in Figure 5. We observe that fine-tuned-CNN outperforms the other models for most of the organisms. Among the 44 selected organisms, fine-tuned-CNN achieves the highest accuracy on 33 of them. Note that all models achieve lower performance classifying tca, ame, and bmo. But even for these organisms, fine-tuned-CNN still obtains a reasonable performance, while for bmo, the other three models perform like a random classifier. Perhaps this performance could be related to the use of miRBase data during the pre-training of fine-tuned-CNN. Nonetheless, this network still outperforms both ML models on previously unseen data such as MirGeneDB and NotBestFold.

**Fig. 5.**
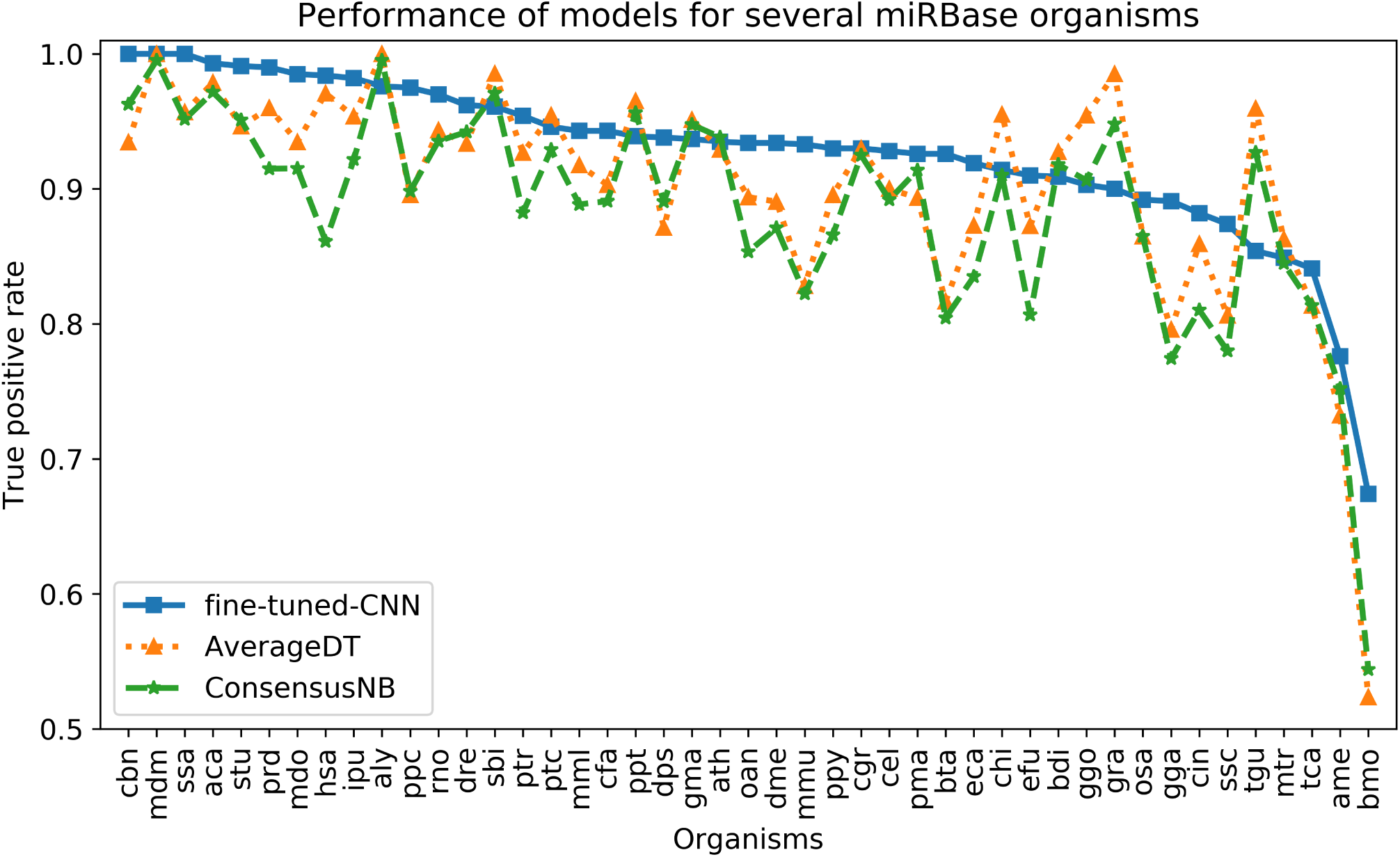
The performances of fine-tuned-CNN and two ML models are evaluated using a subset of miRBase composed of organisms that have a least 200 entries. The plot shows that our CNN outperforms the ensemble models for most of the organisms. For some organisms such as tca, ame, and bmo, pre-miRNAs are difficult to identify. Furthermore, for bmo, both ML models perform similar to a random classifier. In contrast, our CNN is able to learn features for pre-miRNA detection and obtains a performance close to 0.7.

To understand the decision making process of our model in respect to the locations in the hairpin structure, we produce average saliency maps of the true positives and true negatives. Figure 6 is an overlay of both average saliency maps. In this image, pixels relevant for predicting true positives are depicted in green, pixels relevant for predicting non pre-miRNAs are depicted in red, and pixels relevant for predicting both labels are shown in yellow. From a biological point of view these images give us a certain level of sanity check. We see that our model is not sensitive to the hairpin loop that is present in both positive and negative images. This seems to be contrary to the biological understanding that a pre-miRNA must have a terminal loop, however, any RNA structure that folds back on itself must have a terminal loop and, therefore, the majority of negative examples have terminal loops. Thus, this feature is not discriminative for the given data. We also see that the model is particularly sensitive to regions of the image corresponding to G-C bonds, A-U bonds, and G-U bonds. Moreover, it is not sensitive to regions not representing parts of the miRNA molecule such as the edges. This gives us a certain level of confidence that our model behaves as expected, however, we leave a more in-depth analysis of the behavior towards developing explicit interpretations to future work.

**Fig. 6.**
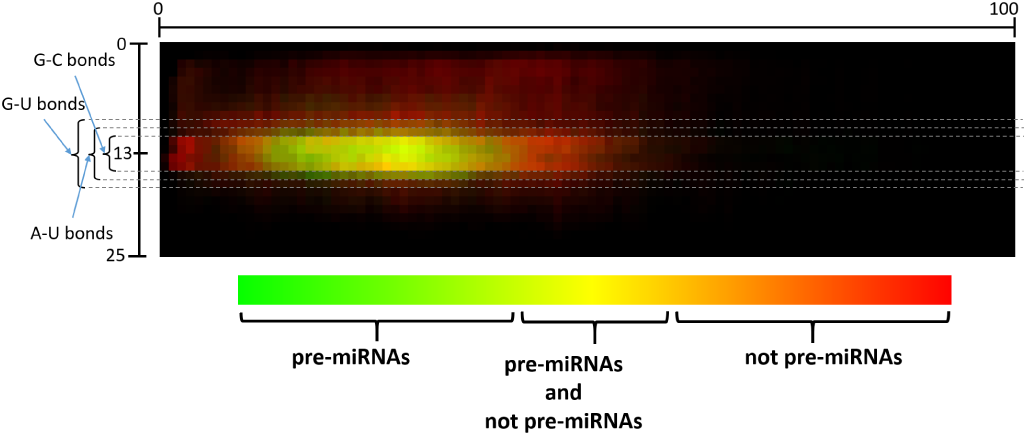
Overlay of average saliency maps of true positives and true negatives computed using the *modhsa* dataset. Relevant regions for pre-miRNA prediction are colored green, and relevant regions for non pre-miRNA prediction are colored red. Moreover, relevant regions for the prediction of both labels are colored yellow. Black pixels indicate irrelevant regions according to the CNN. The regions corresponding to G-C bonds, A-U bonds, and G-U bonds are of particular relevance for the CNN.

**Fig. 7.**
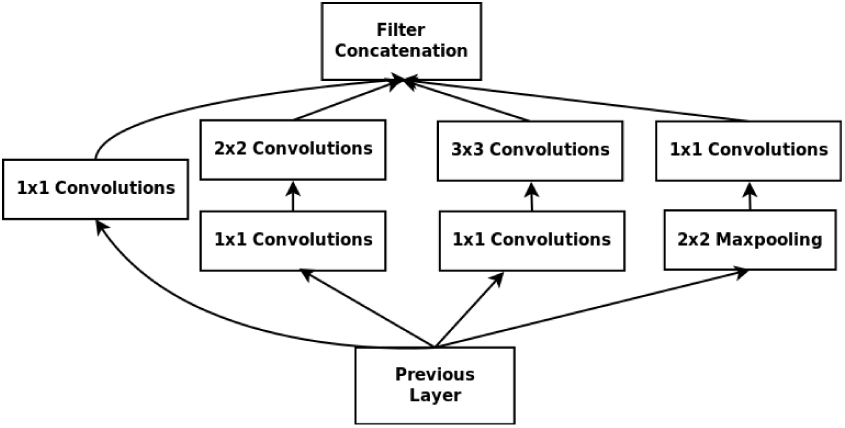
Inception module designed for pre-miRNA detection. It follows a similar structure to the original inception module presented in the GoogLeNet network, but it uses different filter shapes for the convolutional and maxpool layers. The convolutional layers have filters with shape 1×1, 2×2, and 3×3, while the maxpool layer has filter shape of 2×2.

## 4. Discussion

In our experimental study, both proposed CNN models outperform the ensemble models proposed in izMiR, for the NegHsa, NotBestFold, Shuffled, MirGeneDB, mmu, mmu+, and dre+ datasets (Table 2). We achieve similar performance for Pseudo and perform worse only on Chen and Zou. The performance, particularly stands out on NotBestFold dataset, which can be explained by the direct encoding of structure in the images. Sub-optimal folds often do not resemble pre-miRNAs and are, therefore, correctly rejected by both CNNs. On average, fine-tuned-CNN performs better than the other models. Thus, DeepMir uses this model for nucleotide sequence classification.

The results also demonstrate that transfer learning can be used for developing models for other species that have a small number of known pre-miRNAs. In principle, we could select any valid pre-miRNA for the pre-training procedure. Therefore, we should be able to generate CNN models for organisms that have enough pre-miRNAs to perform the fine-tuning procedure. The resulting models could be used on the DeepMir framework, as it allows for the usage of different CNNs. Hence, when necessary, new models could be incorporated to DeepMir.

Besides having excellent performance, one of the main advantage of this approach is that the detection can be done with significantly lower computational complexity than other methods that rely on engineered features. To detect a pre-miRNA sequence we execute two steps; the encoding of the hairpin image and the detection with the neural network. The hairpin encoding algorithm only runs RNAfold once per sequence. All other calculations are significantly simpler. In contrast, several of the ML features are based on statistics and need to rerun RNAfold for the randomly shuffled sequence up to a thousand times with additional feature calculations which may not be trivial (9, 10). For some sequences it may not be possible to create a thousand different sequences. Taken together, some features take a thousand fold longer to be created than an image; and for sequences some additional runtime may be required since a thousand different randomly shuffled sequences cannot be created. The detection with the neural network model is very efficient. The computational complexity of such models scales linearly with the number of neurons and the size of the input. As both of these values are fairly small for today’s computational resources this step is very quick. Furthermore, the execution of the model is highly parallelizable and can be significantly accelerated on suitable hardware.

Overall, the lower computational time required to encode hairpin images makes our pre-miRNA detection framework a feasible choice for even large eukaryotic genomes with millions of putative hairpins.

## 5. Conclusions

MicroRNAs are important in gene regulation and contribute to cell homeostasis. Their dysregulation is a hallmark of disease. However, miRNA and their target detection is experimentally complicated since both are under spatiotemporal control. Therefore, computational methods are important to detect miRNAs and their targets.

We addressed the task of pre-miRNA detection using a deep learning based framework whose key components are a novel hairpin image encoding algorithm and a custom CNN model. To this end, we first encode the primary sequence and secondary structure of nucleotide sequences into RGB images that display spatial patterns and implicitly encode thermodynamics. Then, CNN models are used to classify these images as pre-miRNA or other.

By using this framework, we avoid time consuming feature selection procedures used in ML methods for pre-miRNA detection. When compared against ML pre-miRNA detection methods, our framework outperforms these methods in most of the positive and negative datasets used for evaluation. Therefore, we conclude that this framework can be used for the prediction of novel pre-miRNAs. Additionally, the overlay of average saliency maps show that our CNN model is able to focus on different regions to distinguish between true and false pre-miRNA examples. Further study is required in order to identify how these regions are related to biological structures of pre-miRNAs and whether they are different for different species, classes, or kingdoms.

Even though we achieved state-of-the-art performance, we believe that there may be room for future improvement of both the hairpin encoding algorithm and the CNN models. Specifically, the hairpin algorithm can be extended to incorporate more domain knowledge and accordingly more complex CNN models can be developed for such encodings.

We hope that the exceptional performance of our fine-tuned-CNN model and its applicability for a wide range of species and even for large genomes will enable others to confidently perform pre-miRNA prediction.

## Appendix

### A. CNN architectures

The networks described in Tables 3 to 5 were implemented using the Keras deep learning framework, and their code is available at https://github.com/jacordero/deepmir/. The code includes scripts to design and train VGG-Style, Inception-Style, and ResNet-Style models.

**Table 3.**
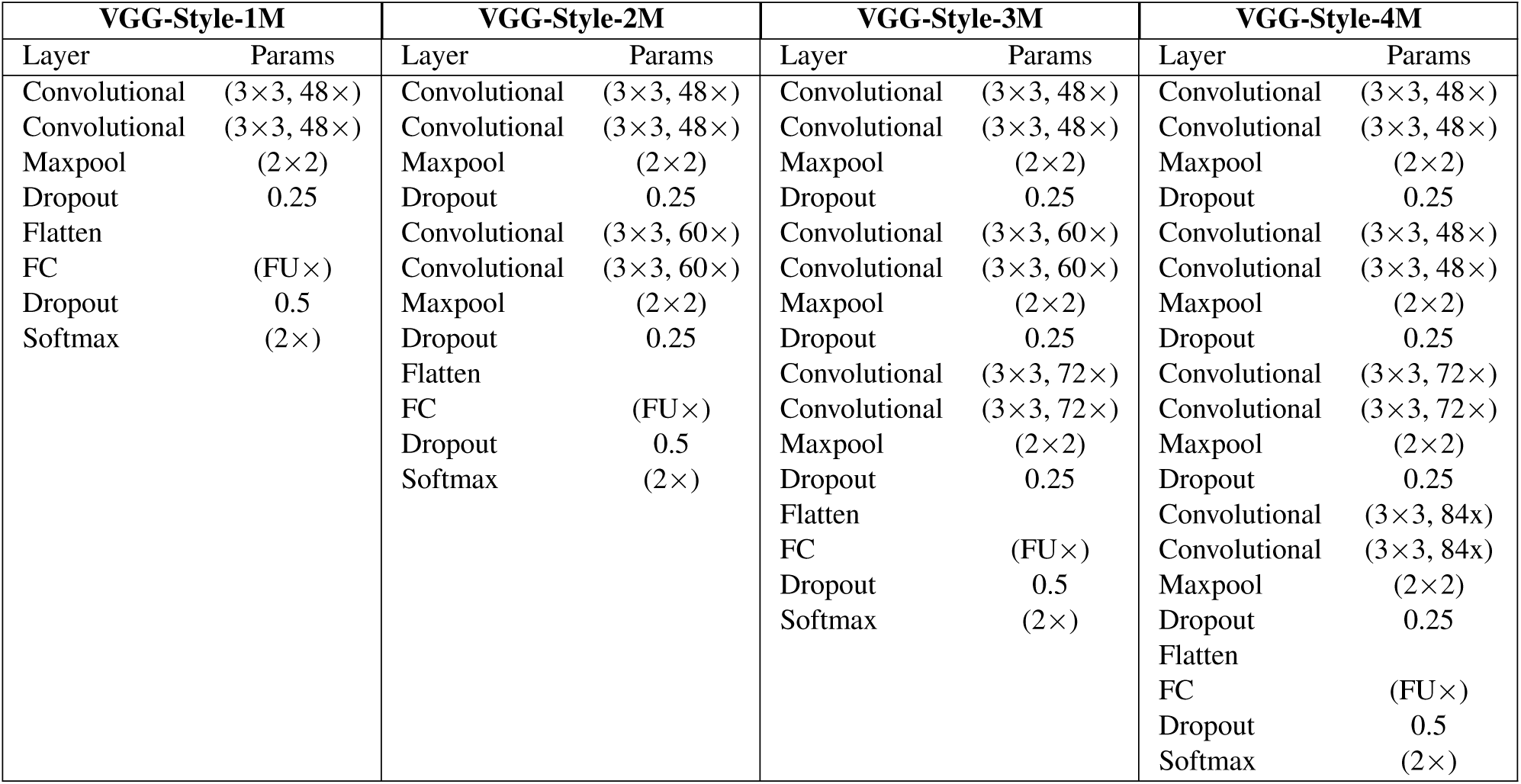
Structure of CNNs that were designed based on the VGG architecture (27). The table shows the structure of VGG-Style networks with up to four convolutional modules (VGG-Style-4M). Each convolutional module is made of two convolutional layers using filter size of 3×3 and the same number of filters (48, 60, 72, or 84) followed by a maxpool layer and a dropout layer. The end of the networks consist of a fully connected module made of a fully connected layer containing a certain number of fully connected units FU (32, 64, 128, or 256) followed by a dropout layer and a softmax layer with two output units (pre-miRNA and non pre-miRNA respectively). In total, we can create 16 different CNNs for a given dataset: four different VGG-Style networks with four different values for the FU parameter.

**Table 4.** Structure of CNNs with up to four modules that were designed based on the Inception architecture (28). The first layer of each network is a convolutional layer with 64 filters of shape 3×3 used to extract low level features. The Inception modules, used to extract spatial patterns at different resolutions, have the structure shown in Figure 7. This Inception module has convolutional layers with different filter shape that the ones presented in the original GoogLeNet, the maxpool layer also differs. All the convolutional layers inside the Inception modules have the same number of filters (CK), which can take four different values: 8, 12, 16, and 20. In total, we can create 16 different CNNs for a given dataset: four different Inception-Style networks with four different values for the CK parameter.

| Inception-Style-1M |  | Inception-Style-2M |  | Inception-Style-3M |  | Inception-Style-4M |  |
| --- | --- | --- | --- | --- | --- | --- | --- |
| Layer/Module | Params | Layer/Module | Params | Layer/Module | Params | Layer/Module | Params |
| Convolutional | $(64 \times, 3 \times 3)$ | Convolutional | $(64 \times, 3 \times 3)$ | Convolutional | $(64 \times, 3 \times 3)$ | Convolutional | $(64 \times, 3 \times 3)$ |
| Inception | $(CK \times)$ | Inception | $(CK \times)$ | Inception | $(CK \times)$ | Inception | $(CK \times)$ |
| Flatten | | Inception | $(CK \times)$ | Inception | $(CK \times)$ | Inception | $(CK \times)$ |
| Softmax | $(2 \times)$ | Flatten | | Inception | $(CK \times)$ | Inception | $(CK \times)$ |
| | | Softmax | $(2 \times)$ | Flatten | | Inception | $(CK \times)$ |
| | | | | Softmax | $(2 \times)$ | Flatten | |
| | | | | | | Softmax | $(2 \times)$ |

**Table 5.** Structure of CNNs with up to four modules that were designed based on the ResNet architecture (35) using fully-preactivated units (36). The first layer of each network is a convolutional layer with 32 filters of shape 3×3 used to extract low level features. The ResNet modules have convolutional layers with a filter shape of 3×3. The number of filters in the convolutional layers is given by the parameter CK that can take four different values: 20, 24, 28, and 32. The GPA layer corresponds to a global average pooling layer that maps 3D feature maps to 1D vectors. The units of such vectors are then connected to the softmax units to produce the output of the network. In total, we can create 16 different CNNs for a given dataset: four different ResNet-Style networks with four different values for the CK parameter.

| ResNet-Style-1M |  | ResNet-Style-2M |  | ResNet-Style-3M |  | ResNet-Style-4M |  |
| --- | --- | --- | --- | --- | --- | --- | --- |
| Layer/Module | Params | Layer/Module | Params | Layer/Module | Params | Layer/Module | Params |
| Convolutional | $(32 \times, 3 \times 3)$ | Convolutional | $(32 \times, 3 \times 3)$ | Convolutional | $(32 \times, 3 \times 3)$ | Convolutional | $(32 \times, 3 \times 3)$ |
| ResNet | $(32 \times, 3 \times 3)$ | ResNet | $(32 \times, 3 \times 3)$ | ResNet | $(32 \times, 3 \times 3)$ | ResNet | $(32 \times, 3 \times 3)$ |
| GPA | | ResNet | $(32 \times, 3 \times 3)$ | ResNet | $(32 \times, 3 \times 3)$ | ResNet | $(32 \times, 3 \times 3)$ |
| Softmax | $(2 \times)$ | GPA | | ResNet | $(32 \times, 3 \times 3)$ | ResNet | $(32 \times, 3 \times 3)$ |
| | | Softmax | $(2 \times)$ | GPA | | ResNet | $(32 \times, 3 \times 3)$ |
| | | | | Softmax | $(2 \times)$ | GPA | |
| | | | | | | Softmax | $(2 \times)$ |

### B. CNNs performance

Table 6 shows the accuracy of the best ten CNN models trained using transfer learning and their corresponding versions without pre-training. Even though 48 different models were trained following the VGG-Style, Inception-Style, and ResNet-Style defined in the tables above, only VGG-Style models belong to the top ten models. The models are sorted according to their average accuracy on both datasets. Tables 7 to 9 show the accuracy of VGG-Style, Inception-Style, and ResNet-Style models respectively. The accuracy of the best models is emphasized in bold.

**Table 6.** Accuracy of top ten pre-trained CNN models including VGG-Style, Inception-Style, and ResNet-Style models sorted by their accuracy. The corresponding non pre-trained models are also included to observe the benefits of pre-training. Only VGG-Style models belong to the top ten CNNs.

| CNN | With pre-training | Without pre-training |
| --- | --- | --- |
| VGG-Style-3M-256FU | **0.9517** | 0.9171 |
| VGG-Style-1M-128FU | 0.9472 | 0.8934 |
| VGG-Style-3M-128FU | 0.9472 | 0.8907 |
| VGG-Style-1M-64FU | 0.9454 | 0.8889 |
| VGG-Style-3M-64FU | 0.9454 | 0.8807 |
| VGG-Style-2M-256FU | 0.9426 | 0.9180 |
| VGG-Style-4M-128FU | 0.9426 | 0.9199 |
| VGG-Style-1M-32FU | 0.9417 | 0.9044 |
| VGG-Style-1M-256FU | 0.9408 | 0.8962 |
| VGG-Style-4M-256FU | 0.9408 | 0.9089 |

**Table 7.** Accuracy of 16 VGG-Style networks trained with and without pre-training. The *modhsa* test dataset is used to evaluate the performance of all trained networks. For pre-trained networks, VGG-Style-3M-256FU has the best performance, while for non pre-trained networks, VGG-Style-2M-128FU and VGG-Style-4M-128FU achieve the best performance.

| CNN | Without pre-training |  |  |  | With pre-training |  |  |  |
| --- | --- | --- | --- | --- | --- | --- | --- | --- |
|  | 32FU | 64FU | 128FU | 256FU | 32FU | 64FU | 128FU | 256FU |
| VGG-Style-1M | 0.9044 | 0.8889 | 0.8934 | 0.8962 | 0.9417 | 0.9454 | 0.9472 | 0.9408 |
| VGG-Style-2M | 0.9162 | 0.9144 | **0.9199** | 0.9180 | 0.9390 | 0.9390 | 0.9381 | 0.9426 |
| VGG-Style-3M | 0.8989 | 0.8807 | 0.8907 | 0.9171 | 0.9362 | 0.9454 | 0.9472 | **0.9517** |
| VGG-Style-4M | 0.9162 | 0.9089 | **0.9199** | 0.9089 | 0.9244 | 0.9381 | 0.9426 | 0.9408 |

**Table 8.** Accuracy of 16 Inception-Style networks trained with and without pre-training. The *modhsa* test dataset is used to evaluate the performance of all trained networks. For pre-trained networks, Inception-Style-2M-8CK has the best performance, while for non pre-trained networks, Inception-Style-2M-16CK achieves the best performance.

| CNN | Without pre-training |  |  |  | With pre-training |  |  |  |
| --- | --- | --- | --- | --- | --- | --- | --- | --- |
|  | 8CK | 12CK | 16CK | 20CK | 8CK | 12CK | 16CK | 20CK |
| Inception-Style-1M | 0.8843 | 0.8880 | 0.5000 | 0.5000 | 0.9311 | 0.9299 | 0.9344 | 0.5000 |
| Inception-Style-2M | 0.8907 | 0.8807 | **0.8953** | 0.8852 | **0.9353** | 0.5000 | 0.9290 | 0.9290 |
| Inception-Style-3M | 0.8834 | 0.8916 | 0.8880 | 0.5000 | 0.9235 | 0.9271 | 0.9335 | 0.9281 |
| Inception-Style-4M | 0.8871 | 0.8843 | 0.8934 | 0.8862 | 0.9281 | 0.9235 | 0.9262 | 0.9226 |

**Table 9.** Accuracy of 16 ResNet networks trained with and without pre-training. The *modhsa* test dataset is used to evaluate the performance of all trained networks. For pre-trained networks, ResNet-Style-1M-24CK has the best performance, while for non pre-trained networks, ResNet-Style-3M-24CK achieves the best performance.

| CNN | Without pre-training |  |  |  | With pre-training |  |  |  |
| --- | --- | --- | --- | --- | --- | --- | --- | --- |
|  | 20CK | 24CK | 28CK | 32CK | 20CK | 24CK | 28CK | 32CK |
| ResNet-Style-1M | 0.5009 | 0.9098 | 0.9044 | 0.9135 | 0.8242 | **0.9208** | 0.5082 | 0.5337 |
| ResNet-Style-2M | 0.5055 | 0.5000 | 0.5000 | 0.5000 | 0.8780 | 0.6366 | 0.6967 | 0.8652 |
| ResNet-Style-3M | 0.5893 | **0.9199** | 0.9044 | 0.5310 | 0.9117 | 0.8470 | 0.6894 | 0.9117 |
| ResNet-Style-4M | 0.9062 | 0.8233 | 0.8925 | 0.9123 | 0.5729 | 0.8525 | 0.5337 | 0.7570 |

### C. Hairpin encoding performance

Here, we present the performance of the hairpin encoding algorithm on a set of 1,000 sequences randomly selected from miRBase. Table 10 shows the time required to generate all hairpin images and Table 11 shows the time that each step of the encoding algorithm requires to generate a single image.

**Table 10.** Running time (in seconds) obtained by creating hairpin images and computing features from a set of 1000 sequences randomly selected from miRBase.

| Run | Hairpin images | Features |
| --- | --- | --- |
| First | 9 | 24,060 |
| Second | 9 | 26,040 |
| Third | 9 | 20,940 |
| Fourth | 9 | 24,060 |
| Fifth | 9 | 24,780 |
| Average | 9 | 23,940 |

**Table 11.** Performance of each step of the hairpin encoding algorithm obtained by encoding a set of 1000 sequences randomly selected from miRBase. The running time of each step is measured in milliseconds. The results show that RNAFold is the most time-consuming operation of the hairpin encoding algorithm. These results agree with the runtime analysis of the hairpin encoding algorithm.

| Step | Minimum | Average | Maximum | Std |
| --- | --- | --- | --- | --- |
| RNAFold | 16 | 29.0098 | 314 | 24.5075 |
| Alignment | 0 | 0.0037 | 2 | 0.07810 |
| Drawing | 0 | 0.2372 | 2 | 0.4283 |
| File writing | 0 | 0.9108 | 5 | 0.5059 |

## References

1. Ayse Elif Erson-Bensan. Introduction to MicroRNAs in Biological Systems, pages 1–14. Humana Press, Totowa, NJ, 2014. ISBN 978-1-62703-748-8. doi: 10.1007/978-1-62703-748-8_1.

2. Yong Huang, Xing-Jia Shen, Quan Zou, Sheng Peng Wang, Shun Ming Tang, and Guo Zheng Zhang. Biological functions of MicroRNAs: A review. Journal of physiology and biochemistry, 67:129–39, 10 2010. doi: 10.1007/s13105-010-0050-6.

3. Hamid Hamzeiy, Rabia Suluyayla, Christoph Brinkrolf, Sebastian Jan Janowski, Ralf Hofestädt, and Jens Allmer. Visualization and Analysis of miRNAs Implicated in Amyotrophic Lateral Sclerosis Within Gene Regulatory Pathways. Studies in health technology and informatics, 253:183–187, 01 2018. doi: 10.3233/978-1-61499-896-9-183.

4. Reinhold Munker and George A. Calin. MicroRNA profiling in cancer. Clinical Science, 121(4):141–158, 2011. ISSN 0143-5221. doi: 10.1042/CS20110005.

5. Enrico De Smaele, Elisabetta Ferretti, and Alberto Gulino. MicroRNAs as biomarkers for CNS cancer and other disorders. Brain Research, 1338:100–111, 2010. ISSN 0006-8993. doi: https://doi.org/10.1016/j.brainres.2010.03.103.

6. Victoria Furer, Jeffrey D. Greenberg, Mukundan Attur, Steven B. Abramson, and Michael H. Pillinger. The role of microRNA in rheumatoid arthritis and other autoimmune diseases. Clinical Immunology, 136(1):1–15, 2010. ISSN 1521-6616. doi: https://doi.org/10.1016/j.clim.2010.02.005.

7. Müşerref Duygu Saçar and Jens Allmer. Machine Learning Methods for MicroRNA Gene Prediction, pages 177–187. Humana Press, Totowa, NJ, 2014. ISBN 978-1-62703-748-8. doi: 10.1007/978-1-62703-748-8_10.

8. Y. Bengio, A. Courville, and P. Vincent. Representation Learning: A Review and New Perspectives. IEEE Transactions on Pattern Analysis and Machine Intelligence, 35(8):1798–1828, Aug 2013. ISSN 0162-8828. doi: 10.1109/TPAMI.2013.50.

9. M. D. Saçar and J. Allmer. Data mining for microrna gene prediction: On the impact of class imbalance and feature number for microrna gene prediction. In 2013 8th International Symposium on Health Informatics and Bioinformatics, pages 1–6, Sep. 2013. doi: 10.1109/HIBIT.2013.6661685.

10. Kwang Loong Stanley Ng and Santosh K. Mishra. De novo SVM classification of precursor microRNAs from genomic pseudo hairpins using global and intrinsic folding measures. Bioinformatics, 23 (11):1321–1330, 2007. doi: 10.1093/bioinformatics/btm026.

11. Chenghai Xue, Fei Li, Tao He, Guo-Ping Liu, Yanda Li, and Xuegong Zhang. Classification of real and pseudo microRNA precursors using local structure-sequence features and support vector machine. BMC Bioinformatics, 6(1):310, Dec 2005. ISSN 1471-2105. doi: 10.1186/1471-2105-6-310.

12. Rukshan Batuwita and Vasile Palade. microPred: effective classification of pre-miRNAs for human miRNA gene prediction. Bioinformatics, 25(8):989–995, 2009. doi: 10.1093/bioinformatics/btp107.

13. W. Khalifa, M. Yousef, M. D. Saçar Demirci, and J Allmer. The impact of feature selection on one and two-class classification performance for plant microRNAs. Peerj, e2135.183–187, 04 2016. doi: 10.7717/peerj.2135.

14. Ivani de ON Lopes, Alexander Schliep, and André CP de LF de Carvalho. The discriminant power of RNA features for pre-miRNA recognition. BMC Bioinformatics, 15(1):124, May 2014. ISSN 1471-2105. doi: 10.1186/1471-2105-15-124.

15. Duygu Saçar, Jan Baumbach, and Jens Allmer. On the performance of pre-microRNA detection algorithms. Nature Communications, 8, Dec 2017. doi: 10.1038/s41467-017-00403-z.

16. Alex Krizhevsky, Ilya Sutskever, and Geoffrey E. Hinton. ImageNet Classification with Deep Convolutional Neural Networks. In Proceedings of the 25th International Conference on Neural Information Processing Systems - Volume 1, NIPS’12, pages 1097–1105, USA, 2012. Curran Associates Inc.

17. Kaiming He, Xiangyu Zhang, Shaoqing Ren, and Jian Sun. Deep Residual Learning for Image Recognition. CoRR, abs/1512.03385, 2015.

18. Ritambhara Singh, Jack Lanchantin, Gabriel Robins, and Yanjun Qi. DeepChrome: deep-learning for predicting gene expression from histone modifications. Bioinformatics, 32(17):i639–i648, 2016. doi: 10.1093/bioinformatics/btw427.

19. Haoyang Zeng, Matthew D. Edwards, Ge Liu, and David K. Gifford. Convolutional neural network architectures for predicting DNA–protein binding. Bioinformatics, 32(12):i121–i127, 2016. doi: 10.1093/bioinformatics/btw255.

20. Ryan Poplin, Pi-Chuan Chang, David Alexander, Scott Schwartz, Thomas Colthurst, Alexander Ku, Dan Newburger, Jojo Dijamco, Nam Nguyen, Pegah Tootoonchi Afshar, Sam Gross, Lizzie Dorfman, Cory Y McLean, and Mark A. DePristo. A universal SNP and small-indel variant caller using deep neural networks. Nature Biotechnology, 36:983–987, 2018.

21. Binh Thanh Do, Vladimir Golkov, Göktuğ Erce Gürel, and Daniel Cremers. Precursor microRNA Identification Using Deep Convolutional Neural Networks. bioRxiv, 2018. doi: 10.1101/414656.10.1101/414656, 16-September-2018, pre-print: not peer reviewed.

22. Adam Gudyś, Michal Wojciech Szcześniak, Marek Sikora, and Izabela Makalowska. HuntMi: an efficient and taxon-specific approach in pre-miRNA identification. BMC Bioinformatics, 14(1):83, Mar 2013. ISSN 1471-2105. doi: 10.1186/1471-2105-14-83.

23. L. Wei, M. Liao, Y. Gao, R. Ji, Z. He, and Q. Zou. Improved and Promising Identification of Human MicroRNAs by Incorporating a High-Quality Negative Set. IEEE/ACM Transactions on Computational Biology and Bioinformatics, 11(1):192–201, Jan 2014. ISSN 1545-5963. doi: 10.1109/TCBB.2013.146.

24. Junjie Chen, Xiaolong Wang, and Bin Liu. iMiRNA-SSF: Improving the Identification of MicroRNA Precursors by Combining Negative Sets with Different Distributions. Scientific Reports, 6:19062, Jan 2016. doi: 10.1038/srep19062.

25. Gary M. Weiss. Mining with Rarity: A Unifying Framework. SIGKDD Explor. Newsl., 6(1):7–19, June 2004. ISSN 1931-0145. doi: 10.1145/1007730.1007734.

26. Ivo L. Hofacker. The Vienna RNA Secondary Structure Server. Nucleic Acids Res, 31:3429–3431, 2003.

27. Karen Simonyan and Andrew Zisserman. Very Deep Convolutional Networks for Large-Scale Image Recognition. CoRR, abs/1409.1556, 2014.

28. C. Szegedy, Wei Liu, Yangqing Jia, P. Sermanet, S. Reed, D. Anguelov, D. Erhan, V. Vanhoucke, and A. Rabinovich. Going deeper with convolutions. In 2015 IEEE Conference on Computer Vision and Pattern Recognition (CVPR), pages 1–9, June 2015. doi: 10.1109/CVPR.2015.7298594.

29. Daniel Smilkov, Nikhil Thorat, Been Kim, Fernanda B. Viégas, and Martin Wattenberg. SmoothGrad: removing noise by adding noise. CoRR, abs/1706.03825, 2017.

30. Jost Tobias Springenberg, Alexey Dosovitskiy, Thomas Brox, and Martin A. Riedmiller. Striving for Simplicity: The All Convolutional Net. CoRR, abs/1412.6806, 2014.

31. François Chollet et al. Keras. https://keras.io, 2015.

32. Diederik P. Kingma and Jimmy Ba. Adam: A Method for Stochastic Optimization. arXiv e-prints, art. 1412.6980, Dec 2014.

33. Isaac Bentwich, Amir Avniel, Yael Karov, Ranit Aharonov, Shlomit Gilad, Omer Barad, Adi Barzilai, Paz Einat, Uri Einav, Eti Meiri, Eilon Sharon, Yael Spector, and Zvi Bentwich. Identification of hundreds of conserved and nonconserved human microRNAs. Nature Genetics, 37(7):766–770, 2005. ISSN 1546-1718. doi: 10.1038/ng1590.

34. William Ritchie, Dadi Gao, and John E. J. Rasko. Defining and providing robust controls for microRNA prediction. Bioinformatics, 28(8):1058–1061, 03 2012. ISSN 1367-4803. doi: 10.1093/bioinformatics/bts114.

35. Kaiming He, Xiangyu Zhang, Shaoqing Ren, and Jian Sun. Deep Residual Learning for Image Recognition. CoRR, abs/1512.03385, 2015.

36. Kaiming He, Xiangyu Zhang, Shaoqing Ren, and Jian Sun. Identity Mappings in Deep Residual Networks. CoRR, abs/1603.05027, 2016.

